# Microeukaryotic indicators of freshwater ecosystem condition from a national river monitoring network

**DOI:** 10.64898/2026.11.16.766975

**Authors:** Amy C. Thorpe, Susheel Bhanu Busi, Jonathan Warren, Kerry Walsh, Daniel S. Read

## Abstract

Microeukaryotes are a diverse and functionally important component of river biofilms, including primary producers, consumers, decomposers, and parasites and symbionts. However, their diversity is likely underestimated, and their composition, environmental drivers, and seasonal dynamics have rarely been studied at large spatial scales. Microbiological monitoring of freshwater ecosystems has historically focused almost entirely on diatoms, so the potential of the wider microbial community as indicators of river condition is unknown. Here, we used 18S rRNA gene metabarcoding of more than 1,600 biofilm samples from 700 sites across England’s River Surveillance Network. We show that communities were dominated by diatoms, chlorophytes, ciliates, and fungi, and shifted from phototroph-dominated biofilms in spring towards more diverse, heterotroph-rich biofilms in autumn. Community turnover was driven by water chemistry rather than geographic distance, with alkalinity, DO, temperature, and nitrate-N among the strongest drivers of composition. Co-occurrence networks resolved assemblages of taxa with shared environmental responses, and threshold indicator taxa analysis identified ASV-level indicators of water chemistry gradients. Community change points along these gradients closely matched those reported for bacteria from the same biofilms, and several taxa declined at lower concentrations than the community as a whole, offering potential early warning of change. These findings show that the microeukaryotic community tracks water chemistry consistently across a national river network and can provide a taxonomically broad set of sensitive and interpretable indicators of river condition that complement diatom-based monitoring and emerging bacterial approaches.

## Introduction

Rivers face increasing pressures from nutrient enrichment, organic and inorganic pollution, land-use change, and climate change (Li et al., 2024; Sayer et al., 2025). Detecting the ecological effects of these pressures requires biological indicators that are sensitive, detect change at management-relevant timescales, and can be applied consistently across large spatial scales (Andrei et al., 2025; Kuehne et al., 2017). River biofilms are particularly well suited to this role. These complex communities of bacteria, archaea, algae, protists, and fungi develop on stones and other substrates, where their taxa respond to the physicochemical conditions of the overlying water (Besemer, 2015). As the biofilm matures, these responses accumulate in its composition, reflecting conditions experienced over time rather than at a single sampling point (Sabater et al., 2026). Biofilm microbiomes are therefore increasingly recognised as indicators that could complement existing biological assessments of river condition (Andrei et al., 2025; Blackman and Altermatt, 2026; Thorpe et al., 2026).

Microeukaryotes are a diverse and functionally important component of river biofilms. They include phototrophs, such as diatoms, chlorophytes, and chrysophytes, which are major primary producers in rivers; consumers, such as ciliates and cercozoans, which graze on bacteria and algae; fungi, which decompose organic matter; and parasites and symbionts of other organisms (Battin et al., 2016; Debroas et al., 2017). These groups interact within and between trophic levels and with bacteria, linking primary production, decomposition, and the transfer of carbon and nutrients through aquatic food webs. Biofilms also serve as a major food source for grazing invertebrates, so changes in microeukaryotic communities can propagate to higher trophic levels (Zancarini et al., 2017).

Despite this diversity, microbiological assessment of freshwater ecosystems has focused almost entirely on diatoms. Diatoms have long been used as indicators of nutrient enrichment and organic pollution (Rimet et al., 2012), and in the UK, are used in routine monitoring through the Trophic Diatom Index (TDI; Kelly and Whitton, 1995). The TDI is based on known sensitivities or tolerances of taxa to environmental pressures and, although originally developed for microscopy, has since been adapted for DNA metabarcoding (Kelly et al., 2020). Other microeukaryotes also show promise for freshwater assessment. For example, combined microscopy and metabarcoding of river biofilms showed that ciliate assemblages differed with hydrological conditions and saprobiological classification, supporting their use as indicators of environmental pressure (Kulaš et al., 2021). Extending assessment beyond diatoms could therefore broaden the biological information obtained from biofilms, and including consumers, decomposers, and parasites alongside primary producers could capture responses across the food web. However, whether these additional groups provide consistent and complementary indicators across a national river network remains unresolved.

Another important challenge is distinguishing responses to anthropogenic pressures from variation along with natural environmental gradients. A national-scale study of diatom metabarcoding found that assemblage-environment relationships, and the ranges over which communities changed, differed among ecoregions, highlighting the importance of environmental context when developing indicators (Smucker et al., 2024). Identifying taxon-specific and community-level change points can show where biological reorganisation occurs along these gradients and whether multiple taxa respond synchronously (Baker and King, 2010). Comparing these responses between bacteria and microeukaryotes could then establish whether different components of the biofilm track the same environmental transitions or provide distinct information. However, beyond diatoms, microeukaryotes have received far less attention than bacteria in environmental microbiology, particularly in rivers, and their diversity is likely to be underestimated (Debroas et al., 2017; Li et al., 2021). As a result, the autecological information needed to develop non-diatom indicators is lacking. Metabarcoding of the 18S rRNA gene can detect this broader community, but its biogeography, environmental drivers, and potential as an indicator of water-quality remain poorly understood.

The River Surveillance Network (RSN) provides a unique opportunity to address these gaps. The RSN is a spatially balanced network of sites designed to assess the condition of England’s rivers, with biofilm samples collected alongside routine water chemistry monitoring. Identifying points along an environmental gradient where taxa and communities change can inform environmental standards and provide early warnings of ecological change (Gsell et al., 2016). Using 16S rRNA gene sequencing of RSN biofilms, we previously identified bacterial indicators with threshold responses along water chemistry gradients, highlighting the potential of microbial indicators for integration into monitoring frameworks (Thorpe et al., 2026; Warren et al., 2026). Metagenomic analysis of the same network further linked biofilm bacterial biogeography and diversity to their functional potential, providing a mechanistic basis for interpreting indicator responses (Thorpe et al., 2025). However, it is not yet known whether microeukaryotes follow the same gradients and thresholds as bacteria, or whether trophic groups respond to water quality in different ways.

Here, we used 18S rRNA gene sequencing to characterise river biofilm microeukaryotic communities at a national scale. Specifically, we aimed to (i) characterise the composition and diversity of biofilm microeukaryotes across England’s river network, (ii) determine the environmental and seasonal gradients that structure these communities, (iii) explore how taxa organise into assemblages with shared responses, and (iv) identify indicator taxa that show threshold responses along water quality gradients.

## Materials and Methods

### Sample collection

Biofilm samples were collected from the Environment Agency’s RSN. Sampling took place in spring (March to May) and autumn (September to November) from 2021 to 2023 (Figure 1), giving 1,643 samples from 700 sites as described in detail in Thorpe et al. (2026). Briefly, biofilm was scraped from submerged stones following Kelly et al. (2020) and stored in DNA preservation buffer. Samples were transported on dry ice to the UK Centre for Ecology & Hydrology (UKCEH), Wallingford, and kept at -20 °C until DNA extraction.

**Figure 1.**
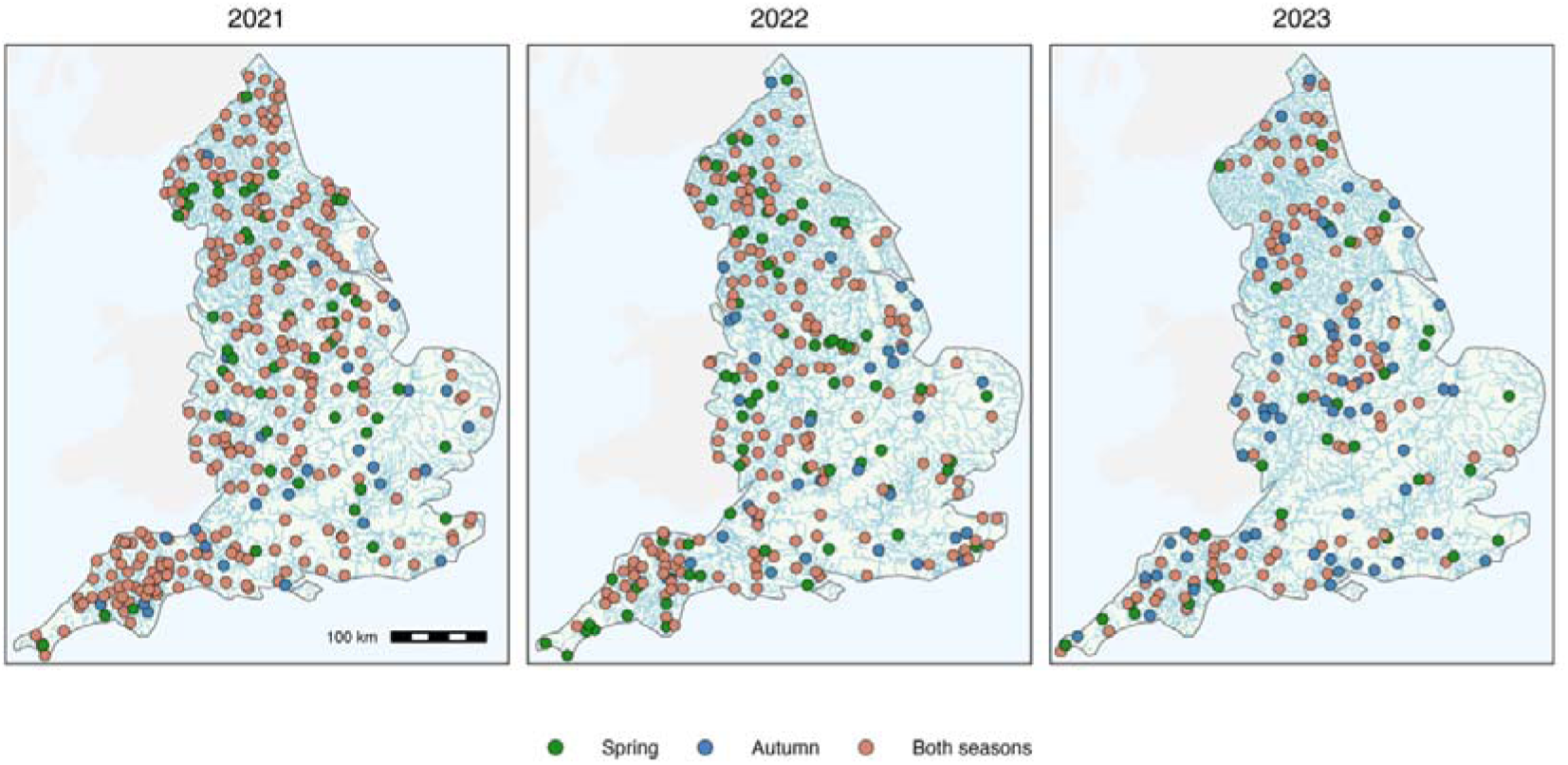
Biofilm sampling sites in rivers across England in 2021-2023. Each panel shows the sites sampled in that year, coloured by whether they were sampled in spring only, autumn only, or both seasons (contains OS data Crown Copyright and database rights, 2026).

### Environmental data collection

Water chemistry for each site was obtained from the Environment Agency’s routine monitoring of surface waters, following the approach of Thorpe et al. (2026), and is publicly available from the Water Quality Archive (Environment Agency, 2026). The variables used were water temperature (°C), pH, conductivity (µS cm ¹), alkalinity (to pH 4.5 as CaCO_3_, mg L^-1^), dissolved oxygen (DO), dissolved organic carbon (DOC), orthophosphate, nitrate-nitrogen (nitrate-N), nitrite-nitrogen (nitrite-N), ammoniacal nitrogen (ammonia-N), and reactive silicon dioxide (SiO_2_), all reported in mg L^-1^. To represent the conditions each biofilm had developed under, rather than a single time point measurement, each variable was averaged over three months up to and including the sampling date, using up to five measurements per sample.

### DNA extraction, PCR amplification, and sequencing

DNA was extracted from 100 µL of biofilm using the Quick-DNA Fecal/Soil Microbe Kit (Zymo Research, CA, U.S.) as described by Thorpe et al. (2026). The 18S rRNA gene libraries were then prepared using a two-step PCR. In the first step, the V4-V5 region of the 18S rRNA gene was amplified with the NSF563 (5’-CGCGGTAATTCCAGCTCCA-3’) and NSR951 (5’-TTGGYRAATGCTTTCGC-3’) primers (Mangot et al., 2013). In the second step, each sample was given a unique combination of forward and reverse indices for demultiplexing (Kozich et al., 2013). Reagents, thermocycling conditions, and the purification, normalisation, and pooling of PCR products are given in the Thorpe et al. (2024) protocol. Pooled libraries were loaded at 1 nM with 30% PhiX and sequenced (paired-end) on an Illumina NextSeq using a P1 600-cycle flow cell. Samples from 2021-2022 and 2023 were sequenced across separate runs.

### Data processing

Sequence processing followed the pipeline described in Thorpe et al. (2026). In brief, reads were demultiplexed and adapters trimmed with the Illumina FASTQ generation pipeline, primers were removed with Cutadapt v4.7 (Martin, 2011), and amplicon sequence variants (ASVs) were inferred with DADA2 v1.26.0 (Callahan et al., 2016). Forward and reverse reads were truncated at 230 and 220 bp, respectively, filtered at a maximum expected error of 2, and merged with an overlap of at least 12 bp. Each sequencing run was processed separately, and the data were then combined before chimeras were removed. Taxonomy was assigned with the naïve Bayesian classifier (Wang et al., 2007) against the Protist Ribosomal Reference database PR² v5.1.1 (Guillou et al., 2012), at a minimum bootstrap confidence of 80.

As this study focuses on microeukaryotes, ASVs from multicellular groups were excluded, mainly metazoans (18% of ASVs) and land plants (Embryophyta, 4%). Potential spurious ASVs with fewer than five reads or found in fewer than five samples were also removed. Each sample was then rarefied to 10,000 reads, the depth at which richness plateaued in most samples. This excluded 284 samples (17%) with too few reads, as well as negative controls.

### Data analysis

ASV counts were converted to relative abundance per sample, and mean relative abundances were calculated across all samples, within seasons, and at the phylum, genus, and trophic level. Trophic assignments were made on ASVs using EukFunc v1.0.0 (Lentendu et al., 2025), which infers functional classes from PR² taxonomy. Functional classes were grouped as phototrophs, mixotrophs, and heterotrophs, with heterotrophs comprising consumers, parasites/symbionts, and saprotrophs. To summarise the trophic balance of each sample, the log_10_ ratio of phototroph to heterotroph relative abundance was calculated. Where EukFunc gave no assignment, unassigned Chlorophyta and Bacillariophyceae were classed as phototrophs, and unassigned Fungi as heterotrophs. All other ASVs without a trophic assignment were left unassigned.

The core community was identified using the microbiome R package v1.30.0 (Lahti and Shetty, 2017), defined as ASVs with a relative abundance above 0.0001 in more than 50% of samples (Thorpe et al., 2026). To describe how the wider community was distributed, the prevalence (percentage of samples in which an ASV was detected) and mean relative abundance were calculated for each ASV.

Alpha diversity, measured as the Shannon index, and observed ASV richness were calculated per sample using the vegan R package v2.7.2 (Oksanen et al., 2025). Many sites were sampled more than once, so differences in diversity and richness between seasons and years were tested with mixed-effects models that included site as a random intercept. Both models included season, year, and their interaction as fixed effects. Shannon diversity was modelled with a linear mixed model while richness was modelled with a negative binomial generalised linear mixed model in lme4 v1.1.37 (Bates et al., 2015).

Community composition was compared using Bray-Curtis dissimilarity on Hellinger-transformed ASV counts. Non-metric multidimensional scaling (NMDS) was performed using the metaMDS() function in vegan. Differences in composition between seasons and years were tested by permutational multivariate analysis of variance (PERMANOVA) using the adonis2() function with 999 permutations on the Bray-Curtis matrix. Terms were tested sequentially (year, season, and year:season interaction) and permutations were restricted within sites to account for repeated sampling. Water chemistry variables were fitted to the ordination as vectors using the envfit() function with 999 permutations. The 20 genera with the highest mean relative abundance across all samples were also fitted to the ordination as weighted averages of sample scores using the wascores() function.

Distance decay was assessed against both geographic and water chemistry distance between samples. Geographic distance was calculated as Haversine distance in km between site coordinates using the geosphere R package v1.5.20 (Hijmans et al., 2016). Water chemistry distance was calculated as the Euclidean distance between samples after standardising each variable to zero mean and unit variance. Community similarity was expressed as 1 – Bray-Curtis dissimilarity. Samples were compared only within the same season and year, giving 137,559 pairwise comparisons. For each season and year, Mantel tests (Spearman, 999 permutations) were used to assess the correlation between community dissimilarity and geographic or water chemistry distance. Partial Mantel tests then assessed each distance while controlling for the other.

Co-occurrence networks were constructed for spring and autumn samples using the microeco R package v2.2.0 (Liu et al., 2021). Within each season, ASVs were retained if they had a mean relative abundance greater than 0.0001 and were present in more than 10% of samples, giving 455 ASVs across 698 spring samples and 511 ASVs across 661 autumn samples. Pairwise associations between ASVs were quantified using Spearman’s rank correlation coefficient (ρ), and only strong and significant positive edges were retained (ρ > 0.3 and false discovery rate (FDR)-adjusted p < 0.01). Network modules were identified with the fast greedy modularity algorithm, and modules of three or fewer ASVs were excluded. Each ASV was assigned a network role from its within-module and among-module connectivity, where module hubs were strongly connected within their own module (within-module connectivity >2.5 and among-module connectivity ≤0.62) and connectors linked different modules (within-module connectivity ≤2.5 and among-module connectivity >0.62) (Liu et al., 2021). Network properties were calculated with the igraph R package v2.2.1 (Csárdi and Nepusz, 2006) and networks were visualised with ggraph v2.2.2 (Pedersen et al., 2025). Within each season, Spearman correlations were computed between module relative abundance and water chemistry variables, with p values adjusted using the Benjamini-Hochberg procedure.

Threshold indicator taxa analysis (TITAN) was used to identify microeukaryotic ASVs that responded to water chemistry gradients, and the points along each gradient where community composition changed most. The analysis followed Thorpe et al. (2026) using the TITAN2 R package v2.4.4 (Baker and King, 2010). In brief, TITAN finds the change point along a gradient that best separates samples with low and high relative abundance of each ASV, based on permutation-standardised indicator values (Z scores). It sums these scores across ASVs to identify community-level change points. The gradients tested were alkalinity, nitrate-N, DO, and water temperature, as analysed for bacteria (Thorpe et al., 2026), which were also among the variables most strongly associated with microeukaryotic community composition as determined with envfit(). TITAN was performed on samples with complete observations for the tested gradients (1,328 samples) and ASVs present in more than 10% of these samples (483 ASVs). Each run was performed with 500 permutations, 500 bootstrap replicates, and a minimum of five samples on either side of a change point. ASVs with purity (consistent direction of response across bootstrap replicates) and reliability (consistent significance) >0.95 were retained as indicators. Those more abundant below the change point were classed as sensitive, declining as the gradient increases, and those more abundant above the change point as tolerant. Community change points are reported from the filtered sum of Z scores (fsumz) based on pure and reliable indicators.

## Results

### Composition, diversity, and seasonal dynamics of river biofilm microeukaryotes

Across the river network, 1,359 biofilm samples from 675 sites (698 spring, 661 autumn) yielded 16,858 microeukaryotic ASVs, spanning 32 phyla, 97 classes, 218 orders, 379 families and 851 genera. Biofilm communities were variable among samples but dominated by a small number of phyla (Figure 2A). Gyrista was the most abundant phylum with a mean relative abundance of 0.34 (± 0.2 SD) across all samples and was the dominant phylum in 40.5% of samples. Gyrista was composed mainly of diatoms (Bacillariophyceae, Coscinodiscophyceae and Mediophyceae; mean = 0.27 ± 0.2). Other dominant phyla were Chlorophyta (mean = 0.27 ± 0.2, dominant in 29.7% of samples), Ciliophora (mean = 0.16 ± 0.2, dominant in 15.2% of samples), and Fungi (mean = 0.14 ± 0.2, dominant in 13.1% of samples).

**Figure 2.**
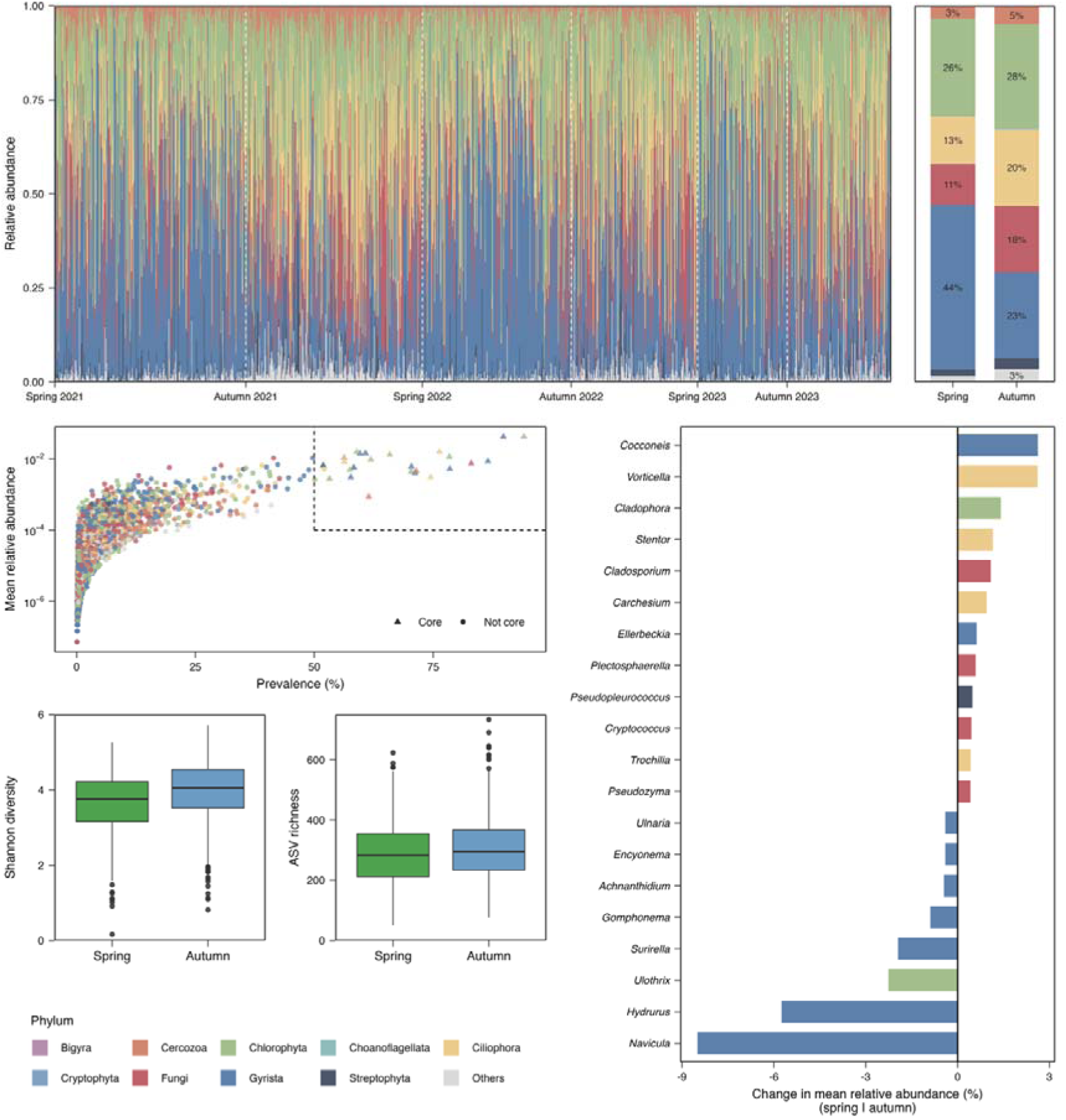
Microeukaryotic community composition of river biofilms. (a) Relative abundance at the phylum level across seasons and years, where phyla with a mean relative abundance <0.001 are grouped as ‘Others’. (b) Phylum-level relative abundance summarised across spring and autumn samples. (c) Genera showing the largest seasonal change in mean relative abundance (autumn – spring, expressed as a percentage). Negative values indicate those more abundant in spring and positive values indicate those more abundant in autumn. (d) Mean relative abundance (log scale) and prevalence of ASVs, where triangles represent core ASVs (relative abundance >0.0001 in >50% of samples, approximated by the dashed box). (e) Shannon diversity, and (f) ASV richness in spring and autumn.

Microeukaryotic composition shifted markedly between seasons, from phototroph-rich biofilms in spring towards a greater contribution of heterotrophs in autumn (Figure 2A, B, and S1). The relative abundance of Gyrista almost halved between spring and autumn (0.44 ± 0.3 vs 0.23 ± 0.2), and the proportion of samples dominated by Gyrista fell from 56.9% to 23.3%. Conversely, Ciliophora (0.13 ± 0.2 vs 0.20 ± 0.2) and Fungi (0.11 ± 0.2 vs 0.18 ± 0.2) each increased by approximately 1.5-fold, while Chlorophyta remained relatively stable (0.26 ± 0.2 vs 0.28 ± 0.2). At the trophic level, phototrophs declined from 0.64 ± 0.2 to 0.51 ± 0.3 and mixotrophs from 0.07 ± 0.2 to 0.01 ± 0.1. Consumers increased from 0.10 ± 0.1 to 0.15 ± 0.1 and parasites/symbionts from 0.03 ± 0.1 to 0.06 ± 0.1 (Figure S1). Seasonal shifts were driven primarily by declines in phototrophic and mixotrophic genera, led by *Navicula*, *Hydrurus*, *Ulothrix*, and *Surirella*. Genera increasing in autumn included the phototrophs *Cocconeis* and *Cladophora*, the ciliates Vorticella and Stentor, and the fungi *Cladosporium* (Figure 2C).

The microeukaryotic community comprised many rare ASVs with low prevalence and relative abundance, with 97.2% of ASVs detected in <10% of samples and 65.7% in <1% (Figure 2D). A small core community of 27 ASVs with a high prevalence and high relative abundance (relative abundance >0.0001 in >50% of samples) accounted for 27.8% of reads. The core community was dominated by Gyrista (10 ASVs) and Chlorophyta (8 ASVs), followed by Ciliophora (5 ASVs), Cercozoa (2 ASVs), Fungi, and Streptophyta (each 1 ASV), and was predominantly phototrophic (18 phototrophs, five consumers, one mixotroph, and one parasite/symbiont ASV). The most prevalent core ASVs were assigned to *Cladophora* (ASV_3, detected in 94.1% of samples), *Navicula* (ASV_2, 89.8%; ASV_27, 86.5%), *Cladosporium* (ASV_32, 83.0%), and *Achnanthidium* (ASV_44, 78.5%) (Figure S2).

Shannon diversity ranged from 0.17 to 5.71 across samples (Figure 2E), and ASV richness ranged from 52 to 733 (Figure 2F). Linear and negative binomial mixed-effects models, with site included as a random effect to account for repeated sampling, showed that both Shannon diversity (spring mean = 3.64 ± 0.8 vs autumn 3.96 ± 0.8; F_1,935_ = 56.08, p < 0.001) and richness (spring mean = 286.2 ± 102.2 vs autumn 305.7 ± 102.4; χ²_1_ = 16.36, p < 0.001) were significantly lower in spring than in autumn. Richness also differed between sampling years (χ²_2_ = 48.45, p < 0.001), while Shannon diversity did not (F_2,1228_ = 2.63, p = 0.07), and there was no significant interaction between season and year for either metric (p > 0.05).

### Environmental drivers of community composition and turnover

Microeukaryotic community composition differed between seasons, with spring and autumn samples separating along NMDS2 (Figure 3A). Season explained 2.47% of the variation in composition (PERMANOVA, F = 34.60, p < 0.001), while year (R² = 0.63%, F = 4.41, p < 0.001) and the season:year interaction (R² = 0.23%, F = 1.58, p < 0.001) were also significant but explained less.

**Figure 3.**
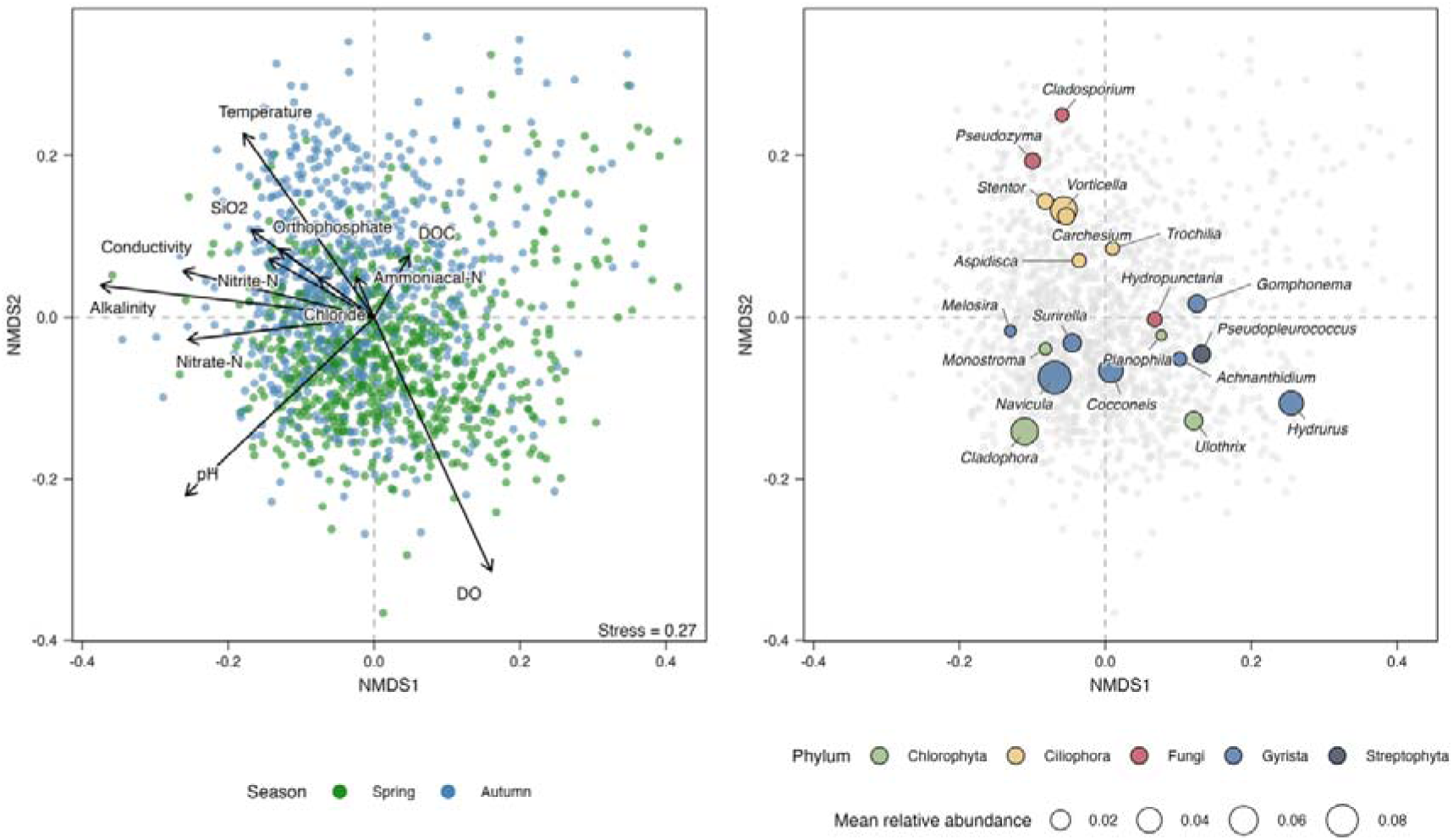
Non-metric multidimensional scaling (NMDS) of a beta diversity Bray-Curtis dissimilarity matrix of river biofilm microeukaryotic communities. (a) Samples coloured by season, with water chemistry vectors fitted. Arrow length is proportional to the correlation. (b) Weighted average positions of the 20 most abundant genera in the ordination space, where points are coloured by phylum and point size is proportional to mean relative abundance.

All tested water chemistry variables were significantly associated with the ordination (p < 0.01; Figure 3A; Table S1). Alkalinity was the strongest (R² = 0.35), followed by DO (0.31), pH (0.28), water temperature (0.20), conductivity (0.18), and nitrate-N (0.16). SiO, nitrite-N, orthophosphate, chloride, DOC, and ammoniacal-N were weaker (<0.10). Alkalinity, conductivity, and nitrate-N aligned with NMDS1, increasing towards negative scores, while DO and temperature aligned more with NMDS2.

The most abundant genera separated by trophic group (Figure 3B). Diatoms (*Navicula*, *Cocconeis*, *Surirella*) and chlorophytes (*Cladophora*, *Ulothrix*, *Monostroma*) were positioned towards negative NMDS2, in line with spring samples and higher DO. In contrast, ciliates (*Stentor*, *Vorticella*, *Carchesium*) and fungi (*Pseudozyma*, *Cladosporium*) were positioned towards positive NMDS2, in line with autumn samples and higher water temperatures. Along NMDS1, *Hydrurus*, *Gomphonema*, and *Pseudopleurococcus* were associated with low alkalinity and low nitrate samples, while *Melosira* and *Cladophora* were associated with high alkalinity and high nitrate samples. The phototroph:heterotroph ratio closely tracked the ordination, mainly along NMDS2 (R² = 0.53, p < 0.001; Figure S4), with more heterotroph-dominated samples associated with positive NMDS2 scores.

Community similarity declined with both geographic and water chemistry distance, but the two relationships differed in strength (Figure 4). Similarity fell from 0.17 between sites less than 25 km apart to 0.13 at 100 km and then stayed between 0.11 and 0.13 up to 600 km (Figure 4A). In contrast, similarity fell steeply with water chemistry distance, from 0.22 between the most chemically similar samples to about 0.09 at distances of 6-9 (Figure 4B). Across each year and season, community dissimilarity was more strongly correlated with water chemistry distance (mean Mantel r = 0.42, range = 0.38-0.48) than with geographic distance (mean r = 0.14, range = 0.10-0.20), and both were significant in every year and season (Table S2). Water chemistry remained a strong predictor after controlling for geography (mean partial r = 0.41, range = 0.36-0.45). In contrast, the geographic correlation almost halved after controlling for water chemistry (mean partial r = 0.08, range = 0.02-0.11). Both relationships were stronger in spring than in autumn (water chemistry r = 0.44 vs 0.40; geography r = 0.16 vs 0.13).

**Figure 4.**
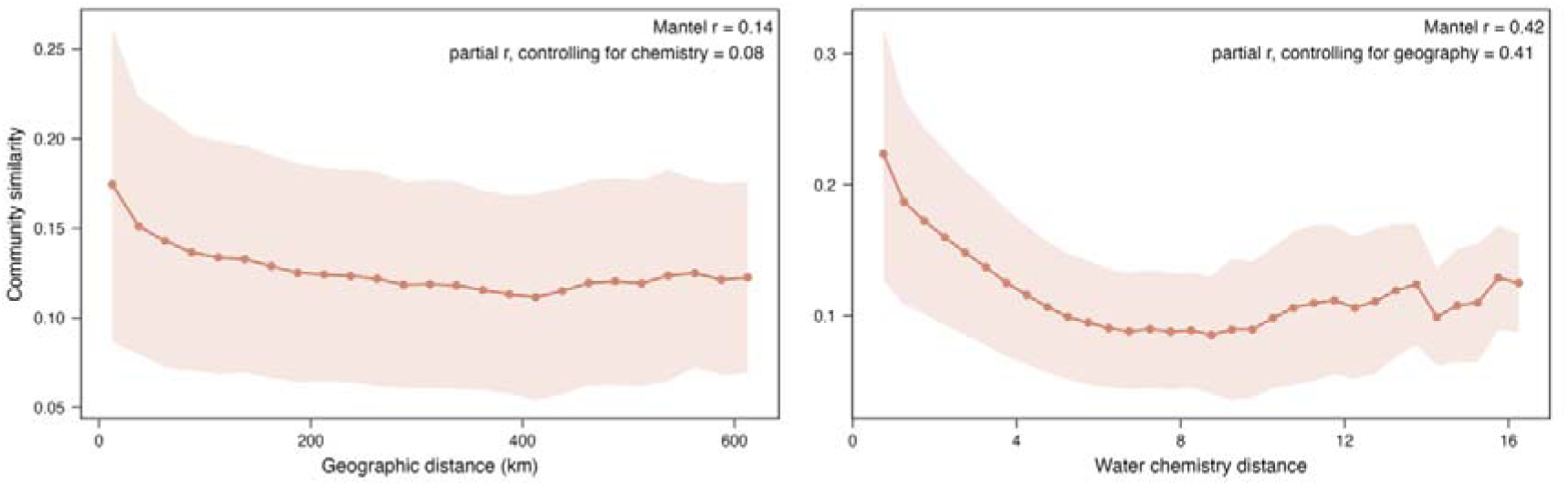
Distance decay of microeukaryotic community similarity with (a) geographic distance between sites, where points show the mean similarity in 25 km distance bins, and (b) water chemistry distance, where points show the mean similarity in 0.5-unit bins. Shaded areas show ± SD. Only pairs of samples from the same season and year were compared. Mean Mantel r and partial Mantel r across each season and year are shown.

### Organisation of seasonal co-occurrence networks

Co-occurrence networks for spring and autumn were highly modular in both seasons but differed in structure and composition (Figure 5A and B). The spring network comprised 376 ASVs connected by 2,406 edges (average degree 12.8). It was more densely connected and clustered than the autumn network (density 0.034 vs 0.022; clustering coefficient 0.51 vs 0.43) and was organised into seven modules (modularity 0.58). The autumn network comprised 404 ASVs and 1,757 edges (average degree 8.7). It was more fragmented, partitioning into ten modules (modularity 0.69). In both seasons, the four largest modules contained the majority of network ASVs (87% in spring, 77% in autumn), and module size dropped sharply after the fourth module (Figure 5E).

**Figure 5.**
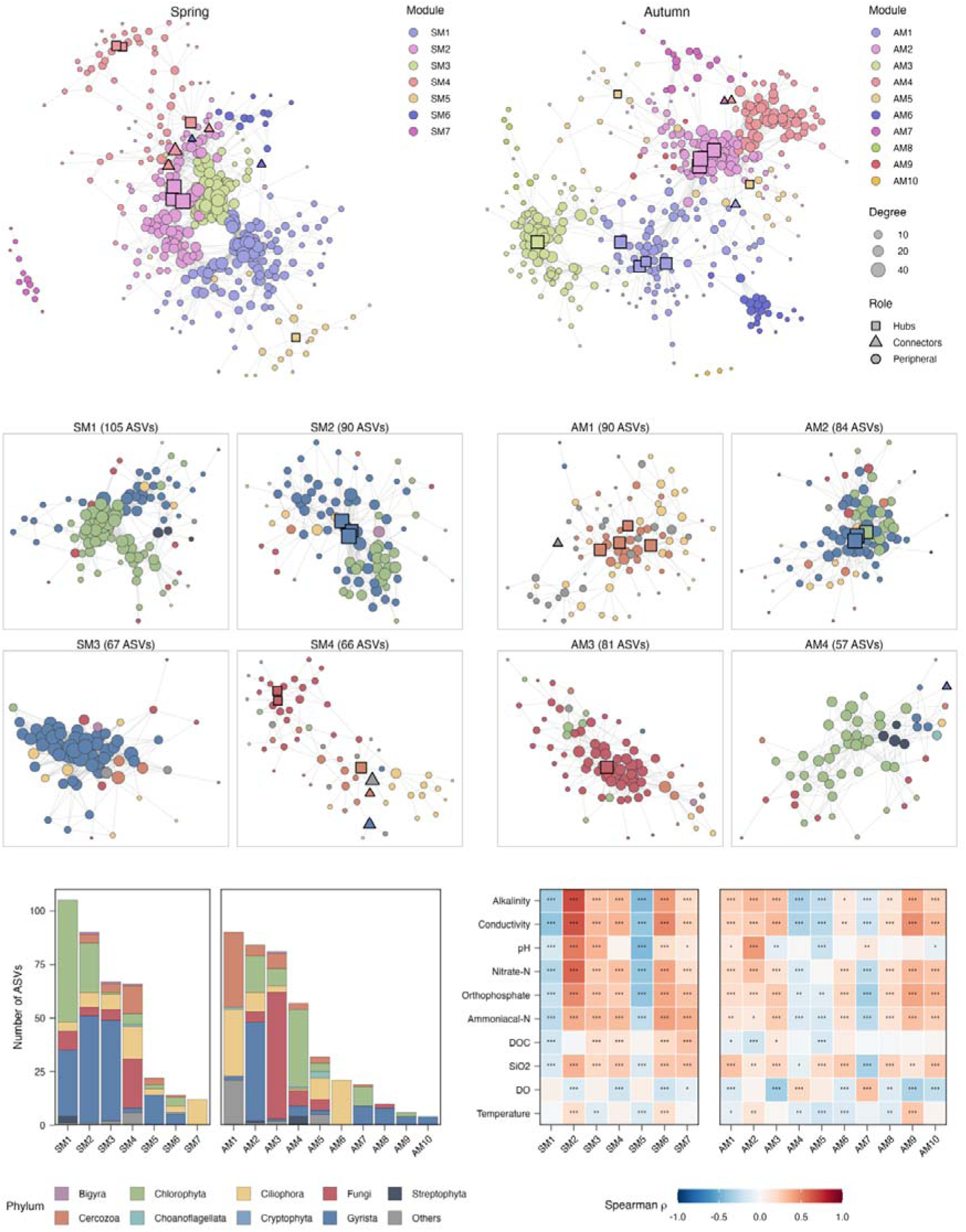
Co-occurrence networks of river biofilm microeukaryotes in (a) spring and (b) autumn. Nodes represent ASVs and are coloured by module (SM, spring modules; AM, autumn modules). Node size is proportional to degree and shape indicates network role (hub, connector, or peripheral node). The four largest modules by number of ASVs in (c) spring and (d) autumn are shown, with nodes coloured by phylum. (e) Number of ASVs and phylum composition in each spring and autumn module. (f) Spearman correlations between module relative abundance and water chemistry, calculated within each season (* p < 0.05, ** p < 0.01, *** p < 0.001).

Module composition reflected seasonal shifts in the community (Figure 5C-E, Figure S6). In spring, the largest modules were dominated by phototrophs. Spring module 1 (SM1, 105 ASVs) and SM2 (90 ASVs) mainly comprised Chlorophyta and Gyrista, but in different proportions (SM1: 54% Chlorophyta and 30% Gyrista ASVs, dominated by *Hydrurus* and *Ulothrix* at the genus level; SM2: 26% Chlorophyta and 56% Gyrista ASVs, dominated by *Cladophora*, *Navicula*, and *Cocconeis*). SM3 (67 ASVs) was diatom-dominated (69% Bacillariophyceae, mainly *Navicula* and *Surirella*). Each of these modules had smaller contributions from Fungi, Ciliophora, and Cercozoa (<10%). SM4 (66 ASVs) was the only large heterotroph module in spring, comprising Fungi (35%), Ciliophora (23%), and Cercozoa (20%), with minor contributions (<10%) from Chlorophyta and Gyrista. Its most abundant genera were the ciliates *Aspidisca* and *Holosticha*. In autumn, two of the four largest modules were dominated by heterotrophs. Autumn module 1 (AM1, 90 ASVs) consisted mainly of Cercozoa (39%) and Ciliophora (34%), and its abundance was dominated by the ciliate *Stentor*. AM3 (81 ASVs) was dominated by Fungi (73%), mainly *Cladosporium* and *Pseudozyma*. In both AM1 and AM3, contribution from Chlorophyta and Gyrista was small (<10%). However, AM2 (84 ASVs) retained the phototroph dominance (55% Gyrista, 20% Chlorophyta), and had a similar genus composition to SM2, dominated by *Cladophora*, *Navicula*, and *Cocconeis*. AM4 (57 ASVs) was mainly Chlorophyta (63%) with a smaller contribution from Fungi (12%) and was dominated by *Pseudopleurococcus* and *Cocconeis*.

The spring network contained seven module hubs and five connectors. The hubs comprised three *Navicula* ASVs in SM2, a *Gomphonema* ASV in SM5, and a *Cladosporium* ASV alongside unassigned Dothideomycetes and Cercozoa ASVs in the heterotroph module SM4. Connectors included *Nitzschia* and *Viridiraptor* ASVs in SM4, and *Stentor* and *Chlamydomonas* ASVs in SM6. The autumn network contained ten module hubs and three connectors. The hubs included four Cercozoa ASVs in AM1, *Cladophora* and two *Navicula* ASVs in AM2, and *Cladosporium* in the fungal module AM3. A *Gomphonema* ASV (AM4) and unassigned Opisthokonta (AM1) and Ulvophyceae ASVs (AM7) comprised the autumn connectors. Two *Navicula* ASVs (ASV_11 and ASV_139) and a *Cladosporium* ASV (ASV_32) were hubs in both seasons.

The strongest associations between ASVs of different genera were mostly between phototrophs, such as the diatoms *Surirella* and *Navicula* (ρ = 0.65 in spring), and Cocconeis and the chlorophytes *Cladophora* and *Monostroma* (ρ = 0.59 in spring and autumn). In autumn, strong associations were observed between fungal ASVs, such as *Cladosporium* and *Alternaria* (ρ = 0.57). Phototroph-heterotroph links made up 13% of edges in spring and 11% in autumn. The strongest were between unassigned Ulvales-relatives and an Endomyxa ASV (ρ = 0.57 in autumn), between the eustigmatophyte *Pseudocharaciopsis* and an unassigned Sanchytriaceae fungus (ρ = 0.57 in autumn), and between *Navicula* and an unassigned chytrid (Rhizophydiales; ρ = 0.48 in spring).

Correlations between modules and water chemistry mostly followed a shared gradient of alkalinity, conductivity, and nutrients, but modules differed in the direction and strength of their responses (Figure 5F). In spring, the two Chlorophyta-Gyrista modules, which differed in genus composition, responded differently to this gradient. SM2 (*Cladophora*, *Navicula*, *Cocconeis*) correlated positively with alkalinity (ρ = 0.67), conductivity (ρ = 0.64), and nitrate-N (ρ = 0.58), while SM1 (*Hydrurus*, *Ulothrix*) correlated negatively with these variables (conductivity ρ = -0.40, alkalinity ρ = -0.34, nitrate-N ρ = -0.29). The diatom-dominated module SM3 (*Navicula*, *Surirella*) and the heterotroph-dominated module SM4 were also positively, but more weakly, associated with this gradient (ρ = 0.25-0.35). Correlations were generally weaker in autumn (maximum |ρ| = 0.46) but followed a similar pattern. The heterotroph-dominated modules AM1 and AM3 correlated positively with most measured variables and negatively with DO, most strongly for the fungal module AM3 (DO ρ = -0.35). As in spring, the two phototroph-dominated modules diverged. AM2, which resembled SM2 in genus composition, correlated positively with pH (ρ = 0.44), alkalinity (ρ = 0.32), and nitrate-N (ρ = 0.30). AM4 correlated negatively with alkalinity (ρ = -0.32), conductivity (ρ = -0.31), and nitrate-N (ρ = -0.15).

### Threshold responses of microeukaryotic indicators to water chemistry gradients

TITAN identified 469 of the 483 tested ASVs (97%) as pure and reliable indicators of at least one gradient (Figure 6). Many ASVs responded to multiple gradients, with 165 (35%) indicators of all four gradients and only 37 (8%) of a single gradient (Figure S7). Indicators also included most key community members, with 26 of the 27 core ASVs identified as indicators, and 89% of indicators (416 ASVs) were network members, including all module hubs and connectors.

**Figure 6.**
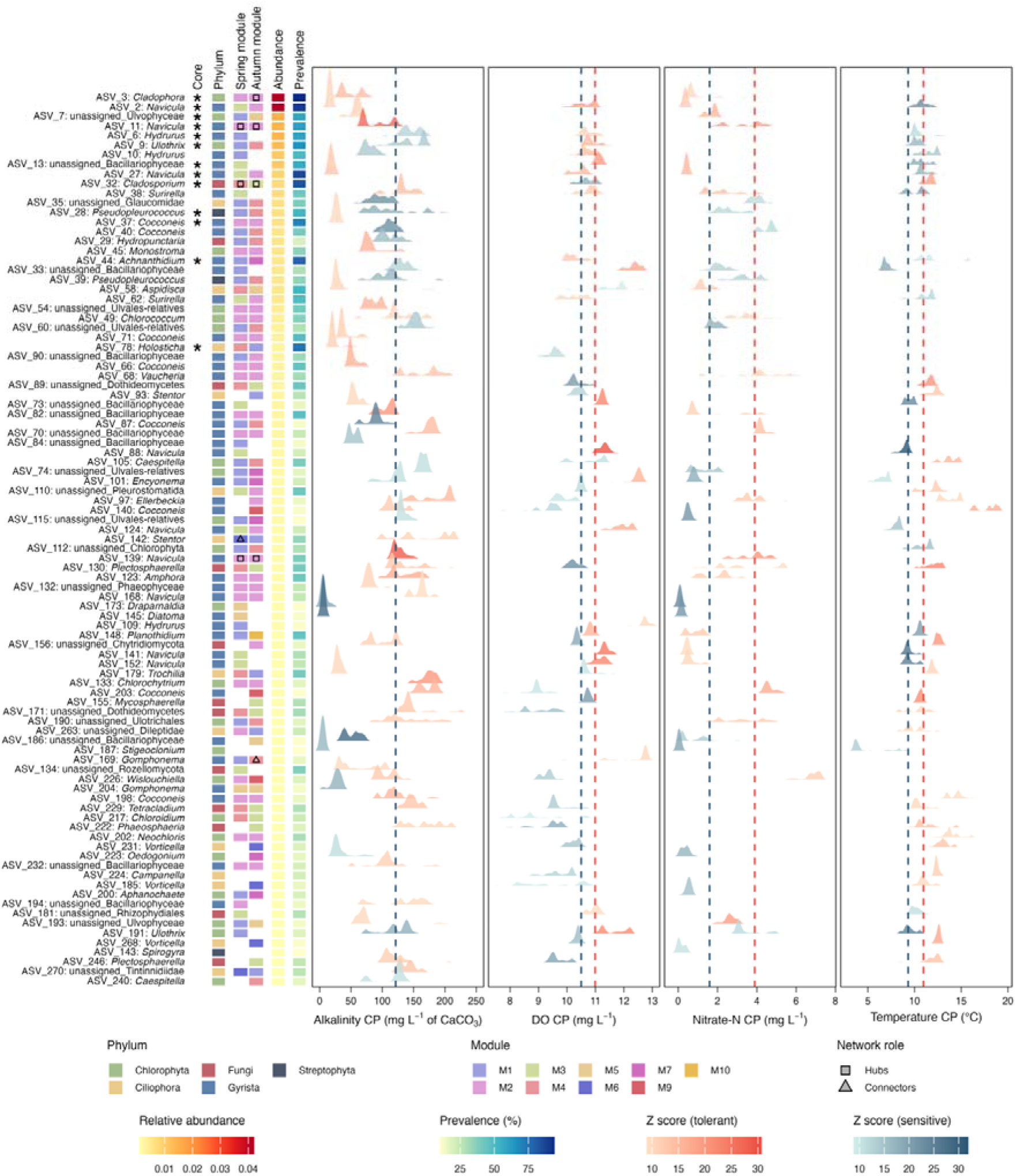
Threshold indicator taxa analysis (TITAN) of microeukaryotic ASVs responding to (a) alkalinity, (b) DO, (c) nitrate-N, and (d) temperature. Ridges represent the distribution of bootstrap change points for each ASV, colour gradients correspond to the magnitude of the response at the median change point (Z score) for individual tolerant and sensitive ASVs, and dashed lines show the community-level change point (CP) for all tolerant (red) and sensitive (blue) ASVs. Core ASVs are marked with stars. The phylum, network role, mean relative abundance and prevalence (%) are shown for each ASV. Each ASV is labelled by genus, and where the genus is unassigned, ASVs are labelled by the most specific assigned taxonomic level. Only ASVs with TITAN purity and reliability scores >0.95, Z scores >10 and mean relative abundance >0.001 are shown.

Alkalinity had the most indicators (369 ASVs; 288 tolerant and 81 sensitive), and tolerant and sensitive ASVs shared a community change point at 121.4 mg L ¹ CaCO_3_ (Figure 6A). DO had 348 indicators (149 tolerant, 199 sensitive), with community change points at 10.99 mg L ¹ for tolerant and 10.50 mg L ¹ for sensitive ASVs (Figure 6B). Nitrate-N had 321 indicators (237 tolerant, 84 sensitive), with change points at 3.89 and 1.60 mg L ¹ (Figure 6C), and temperature had 341 indicators (198 tolerant, 143 sensitive), with change points at 10.95 and 9.32 °C (Figure 6D).

Individual change points varied widely around the community change points, and some ASVs responded before or after the community (90% bootstrap intervals beyond the community change point). We focus on ASVs that responded before the community in the direction of deterioration, at lower values along the alkalinity, nitrate-N, and temperature gradients and at higher values along the DO gradient. Along the alkalinity and nitrate-N gradients, 28 and 21 sensitive ASVs declined below the community change point. The strongest of these, which declined along both gradients, were the chlorophytes *Draparnaldia* (ASV_173) and *Stigeoclonium* (ASV_187) and the diatom *Diatoma* (ASV_145), at 5-6 mg L ¹ CaCO_3_ and 0.03-0.15 mg L ¹ nitrate-N. Similarly, 48 and 49 tolerant ASVs increased below the community change point. These included *Cladophora* (ASV_3; 42 mg L ¹ CaCO_3_ and 0.65 mg L ¹ nitrate-N), *Cocconeis* (ASV_37; 27 mg L ¹ CaCO_3_ and 0.27 mg L ¹ nitrate-N), and *Navicula* (ASV_2 and ASV_27; 16-18 mg L ¹ CaCO_3_ and ∼0.4 mg L ¹ nitrate-N). Along the DO gradient, 33 tolerant ASVs had change points above the community change point. These were mainly diatoms and chlorophytes such as *Navicula* (ASV_88 and ASV_141; 11.3 mg L ¹) and the chrysophyte *Hydrurus* (ASV_109; 12.9 mg L ¹). A further 11 sensitive ASVs increased above the community change point, but all had weak responses or low abundance (Z score < 10 or mean relative abundance < 0.001). Temperature had few early responders. Only 11 sensitive ASVs declined below the community change point, including *Navicula* (ASV_124, 8.3 °C) and an unassigned diatom (ASV_33, 6.8 °C), which were also early responders to DO. Three tolerant ASVs increased below the community change point, all with weak responses or low abundance.

## Discussion

Microeukaryotes are a diverse but understudied component of freshwater ecosystems (Debroas et al., 2017; Li et al., 2021), and while diatoms are widely used as freshwater indicators (Kelly et al., 2020), the potential of the wider community as indicators of water quality has not previously been assessed at a national scale. Using biofilms from across England’s river network, we show that microeukaryotic communities are structured by season and water chemistry gradients, organise into distinct assemblages of taxa with shared responses, and exhibit threshold responses relevant to monitoring. Together, these findings demonstrate that microeukaryotes can provide sensitive, interpretable indicators of river condition that complement diatom-based and emerging bacterial approaches.

A clear seasonal shift from phototroph-dominated biofilms in spring towards more heterotroph-dominated biofilms in autumn was observed across the river network. This shift was driven mainly by a decline in diatoms, with Gyrista almost halving between spring and autumn, while the proportion of Chlorophyta remained more stable across seasons. Seasonal reorganisation also occurred within the phototroph community, with *Navicula*, *Surirella*, *Hydrurus*, and *Ulothrix* more abundant in spring, and *Cladophora* and *Cocconeis* more abundant in autumn. The dominance of diatoms in spring is consistent with the spring diatom bloom reported in temperate rivers (Muhammad et al., 2021; Rose et al., 2021), when cold water and increasing day length before canopy closure, combined with the delivery of nutrients over winter, favour rapid diatom growth (Wang et al., 2022).

Much of the autumn increase in heterotrophs was driven by fungi such as *Cladosporium*, *Alternaria*, and *Pseudozyma*, which are not typically aquatic fungi and are more commonly found on plant material and in soil (Bensch et al., 2012; Kitamoto, 2019; Thomma, 2003). Their increase coincides with leaf fall and higher runoff in autumn, so they may indicate inputs of terrestrial organic matter to rivers rather than resident biofilm populations (Matsuoka et al., 2019; Miura et al., 2026). Peritrich and heterotrich ciliates, including *Vorticella*, *Carchesium*, and *Stentor*, also increased in autumn. These are mostly attached, or sessile, filter feeders that consume bacteria and fine particles (Becz and Török, 2025), and their increase may track higher bacterial production in a mature autumn biofilm with more organic matter, including from decomposing leaf litter. Autumn biofilms were also more diverse and richer in ASVs than spring biofilms, although the difference was modest. This is consistent with an older autumn biofilm being colonised by heterotrophs and receiving additional taxa from terrestrial sources. In contrast, spring biofilms often re-establish after winter high flows and may be dominated by fast-growing phototrophs (Davie et al., 2012).

Community turnover was better explained by water chemistry than by geographic distance. After controlling for geography, the correlation with water chemistry remained strong, while the correlation with geographic distance almost disappeared; similarity declined with geographic distance only over the first ∼100 km. This suggests that, at the scale of England, microeukaryotic communities are assembled mainly by environmental filtering or species sorting, with spatial structure limited to the greater similarity of nearby sites. Rivers are highly connected systems, in which downstream flow can transport cells between sites within the same catchment (Zhang et al., 2023), which may explain this local similarity. Beyond this scale, the weak geographic signal is consistent with the wide dispersal of many microeukaryotes (Finlay, 2002).

Alkalinity was the variable most strongly associated with community composition, as it was for bacteria (Thorpe et al., 2026). Alkalinity, conductivity, and nitrate-N aligned with the same ordination axis and were positively correlated across these sites (Thorpe et al., 2026). Catchment geology largely determines alkalinity and conductivity, which are lower in rivers draining siliceous rocks and higher in rivers draining calcareous rocks (Kney and Brandes, 2007), while nitrate-N mainly reflects inputs from agriculture and wastewater (Li et al., 2023). Across England these coincide, as arable and urban land are concentrated in lowland, calcareous catchments (Thorpe et al., 2025), so the axis represents a combined gradient from low alkalinity (soft), nutrient-poor rivers to high alkalinity (hard), nutrient-enriched rivers. DO and water temperature aligned mainly with the seasonal separation of samples and the phototroph:heterotroph ratio. DO is likely to be partly a result of biofilm activity as well as a driver, with higher DO in spring reflecting colder water and photosynthesis by phototroph-rich biofilms, and lower DO in autumn reflecting warmer water and greater rates of respiration (Halliday et al., 2016).

The relationship between community composition and water chemistry was slightly stronger in spring than in autumn. Winter high flows can scour and reset biofilms (Kendrick and Huryn, 2015), so spring communities may reassemble under strong, shared environmental filters. Autumn biofilms, in contrast, have developed over a longer period and receive more variable inputs, such as leaf litter and its associated terrestrial fungi, which may weaken the water chemistry signal. The spring network was also denser and less fragmented than the autumn network, consistent with stronger environmental sorting in spring. Although these patterns were consistent, season and water chemistry each explained a modest share of the variation in composition. Factors not measured in this study are likely to explain part of the remaining variation, including light availability and riparian shading (Bengtsson et al., 2018; Halliday et al., 2016), flow history (Hope et al., 2025), biofilm age (Brasell et al., 2015; Lyautey et al., 2005), and grazing by invertebrates (Lv et al., 2024).

The co-occurrence networks are likely to represent groups of taxa that respond to the environment in the same way and share habitat preferences, although some associations may also reflect interactions. Module hubs included several *Navicula*, *Cladophora*, and *Cladosporium* ASVs, some of which were hubs in both seasons, and *Cladophora* and *Cladosporium* were also among the most prevalent core ASVs. Hubs are therefore likely to be abundant, widespread taxa that co-occur with many others, so their central position may reflect a broad distribution rather than a keystone role in the community (Zamkovaya et al., 2021).

In spring, SM1 and SM2 were both composed mainly of Chlorophyta and Gyrista but differed in their dominant genera and responded in opposite directions to water chemistry gradients. SM1 was dominated by *Hydrurus* and *Ulothrix*. *Hydrurus* is a chrysophyte of cold, fast-flowing water (Rakonjac and Simić, 2024), and members of *Ulothrix* also tolerate or prefer cold conditions (Vadeboncoeur et al., 2021). SM2 was dominated by *Cladophora*, *Navicula*, and *Cocconeis*. *Cladophora* is typical of high alkalinity, nutrient-enriched rivers (Dodds and Gudder, 1992), and *Cocconeis* and many *Navicula* species tolerate nutrient enrichment (Kelly and Witton, 1995). *Cocconeis* was strongly associated with *Cladophora*, and this assemblage persisted across seasons, dominating both SM2 and AM2, which had a similar composition and a similar response to water chemistry. *Cocconeis* is a well-known epiphyte that attaches to *Cladophora* and other filamentous algae (Marks and Power, 2001). This association may therefore reflect a physical host-epiphyte relationship as well as a shared preference for high alkalinity nutrient-enriched rivers. The persistence of this assemblage across seasons suggests that it is a stable feature of biofilms in enriched rivers, while the *Hydrurus*-*Ulothrix* assemblage was restricted to spring.

Links between phototrophs and heterotrophs made up a small share of associations (13% in spring and 11% in autumn), but several of the strongest involved heterotrophs from lineages known to parasitise or feed on algae. *Navicula* was associated with an unassigned chytrid of the Rhizophydiales, an order that includes parasites of diatoms and other algae (Letcher et al., 2006). Ulvales were associated with an Endomyxa ASV, a group that includes vampyrellid amoebae that feed on green algae (Hess and Suthaus, 2022), and the eustigmatophyte *Pseudocharaciopsis* was associated with a member of the Sanchytriaceae, a family of chytrid-like fungi whose described members parasitise algae (Galindo et al., 2021).

Community change points for sensitive and tolerant microeukaryotes closely matched those identified for bacteria within the same biofilms for each gradient tested (Thorpe et al., 2026). This suggests that bacteria and microeukaryotic communities may reorganise at similar points along these gradients and reflect shared responses of both domains to the same conditions. It may also reflect the close coupling of bacteria and microeukaryotes within biofilms, where phototrophs supply organic carbon to heterotrophic bacteria, and protists graze on bacteria (Zancarini et al., 2017). However, both analyses used the same samples and water chemistry data, and change points could therefore only fall within the observed values, which may contribute to the agreement. Further studies across a wider range of river types and water chemistry, ideally with independent samples and experimental manipulation of key gradients, are needed to test whether these thresholds hold more widely.

Individual ASV change points revealed responses before the corresponding community change points. For example, the chlorophytes *Draparnaldia* and *Stigeoclonium* and the diatom *Diatoma* declined at nitrate-N concentrations of 0.03-0.15 mg L ¹, well below the community change point of 1.60 mg L ¹ for sensitive taxa. Conversely, *Cladophora*, *Cocconeis*, and *Navicula* ASVs increased at 0.27-0.65 mg L ¹ nitrate-N, well below the tolerant change point of 3.89 mg L ¹. Along the DO gradient, *Navicula* and *Hydrurus* ASVs declined below 11.3-12.9 mg L ¹, above the community change point of 10.99 mg L ¹ and would therefore be expected to be lost first as DO falls, while temperature had few early responders. These responses illustrate the value of examining individual taxa alongside aggregate community change, because particular taxa may respond outside the range of greatest collective change (King and Baker, 2010). These ASVs are therefore candidates for detecting biological change before the wider community responds. However, their positions along the gradient do not show that they provide advance warning of deterioration through time; repeated observations at individual sites are needed to determine whether their responses precede broader community change. Interpretation also depends on the gradient considered. TITAN identifies taxa that increase or decrease with an environmental variable, but these directions do not themselves indicate ecological improvement or deterioration. For example, a decline with increasing alkalinity may identify taxa associated with naturally soft-water rivers, and an increase with DO does not imply tolerance of degraded conditions. As assemblage-environment relationships also differ regionally (Smucker et al., 2024), the change points reported here should be regarded as locations of compositional reorganisation within the sampled river network, rather than validated boundaries between ecological condition classes.

The core community was dominated by phototrophs, including many of the diatoms already targeted by current monitoring, while most heterotroph diversity sat in the rarer, more variable part of the community. Microeukaryotes therefore add information beyond diatoms, and co-responding taxa, such as the *Cladophora*-*Cocconeis* and *Hydrurus*-*Ulothrix* assemblages, may provide more robust indicators than single taxa. Chlorophytes, ciliates, fungi, and other non-diatom groups further broadened the taxonomic and trophic range of candidate indicators, consistent with evidence for ciliate-based assessment of rivers (Kulaš et al., 2021). However, comparisons with existing diatom assessments and independent measures of ecological condition are needed to establish whether additional taxa, co-responding assemblages, or combined bacterial-microeukaryotic indicators improve ecological assessment. Routine application will also require validation across independent catchments and years, accounting for river type, season, and natural differences in alkalinity, particularly as diatom-environment relationships vary regionally (Smucker et al., 2024). In addition, TITAN tested only 483 prevalent ASVs of the 16,858 detected, leaving the indicator potential of rarer taxa unresolved. Finally, widespread responses to environmental gradients do not establish diagnostic specificity, so operational indicators must show repeatable responses that distinguish environmental pressures from natural variation.

Additional limitations of our study must also be noted for future studies or applications within monitoring schemes. The number of 18S rRNA gene copies per cell varies by orders of magnitude among microeukaryotes and is particularly high in ciliates, so relative abundances do not equate to cell numbers or biomass (Gong et al., 2013; Zhu et al., 2005). This mainly affects comparisons between taxa and has less effect on comparisons of the same ASV across samples, which underlie the co-occurrence networks and threshold indicator analysis. The primer pair used here was selected in silico for best recovery of freshwater microeukaryotes (Mangot et al., 2013), though this was based on planktonic rather than biofilm communities, and some benthic lineages may therefore be underrepresented. Reference databases such as PR^2^ are incomplete and many ASVs (51%) could not be assigned to genus. Trophic assignments from EukFunc were developed mainly for soil organisms and are coarse for freshwater taxa (Lentendu et al., 2025), and many ASVs had no trophic assignment (23%). Furthermore, the water chemistry variables were strongly correlated with each other (Thorpe et al., 2026), and so the responses described here identify associations with water chemistry gradients rather than the effects of individual pressures.

To conclude, river biofilm microeukaryotes across England’s river network were structured by gradients of water chemistry, particularly alkalinity and nitrate-N, and by a seasonal shift from phototroph-dominated to more heterotroph-rich biofilms. Communities were assembled mainly by these water chemistry gradients rather than geographical distance. Assemblages of taxa with shared responses formed and taxa showed threshold responses that closely matched those of bacteria from the same samples. Taxa beyond diatoms, particularly chlorophytes, ciliates, and fungi, responded clearly to water quality, and several show change points before the community-wide threshold, supporting the inclusion of the wider microeukaryotic community in freshwater ecosystem monitoring. Further work is required to follow seasonal succession at a higher temporal resolution, to extend this approach to a wider range of river types with distinct water chemistry gradients, to test responses through experimental manipulation, and to use targeted markers for poorly resolved groups.

## Data availability

The R scripts to process and analyse the data are available at: https://github.com/amycthorpe/biofilm_18S_analysis. Raw sequence reads have been deposited in the European Nucleotide Archive (ENA) at EMBL-EBI under accession number PRJEB129764 available at: https://www.ebi.ac.uk/ena/browser/view/PRJEB129764. All the data presented are available on Zenodo at: https://doi.org/10.5281/zenodo.24136801.

## Author contributions

**ACT**: conceptualisation, methodology, data curation, investigation, formal analysis, visualisation, writing – original draft, writing – review and editing. **SBB**: conceptualisation, methodology, writing – review and editing. **JW**: conceptualisation, funding acquisition, writing – review and editing. **KW**: conceptualisation, funding acquisition, writing – review and editing. **DSR**: conceptualisation, methodology, funding acquisition, writing – review and editing.

## Acknowledgments

Thank you to Dr Jonathan Porter, Sean Butler, and Alan Wan from the Environment Agency National Laboratory Service for their support collating and shipping the biofilm samples for further processing. This work was funded by the Environment Agency under research project SC220034. ACT, SBB, and DSR were supported by Natural Environment Research Council (NERC) grant NE/X015947/1 and NE/Z000173/1. KW and JW were supported by NERC grant NE/X015777/1.

## Supplementary Information

**Figure S1.**
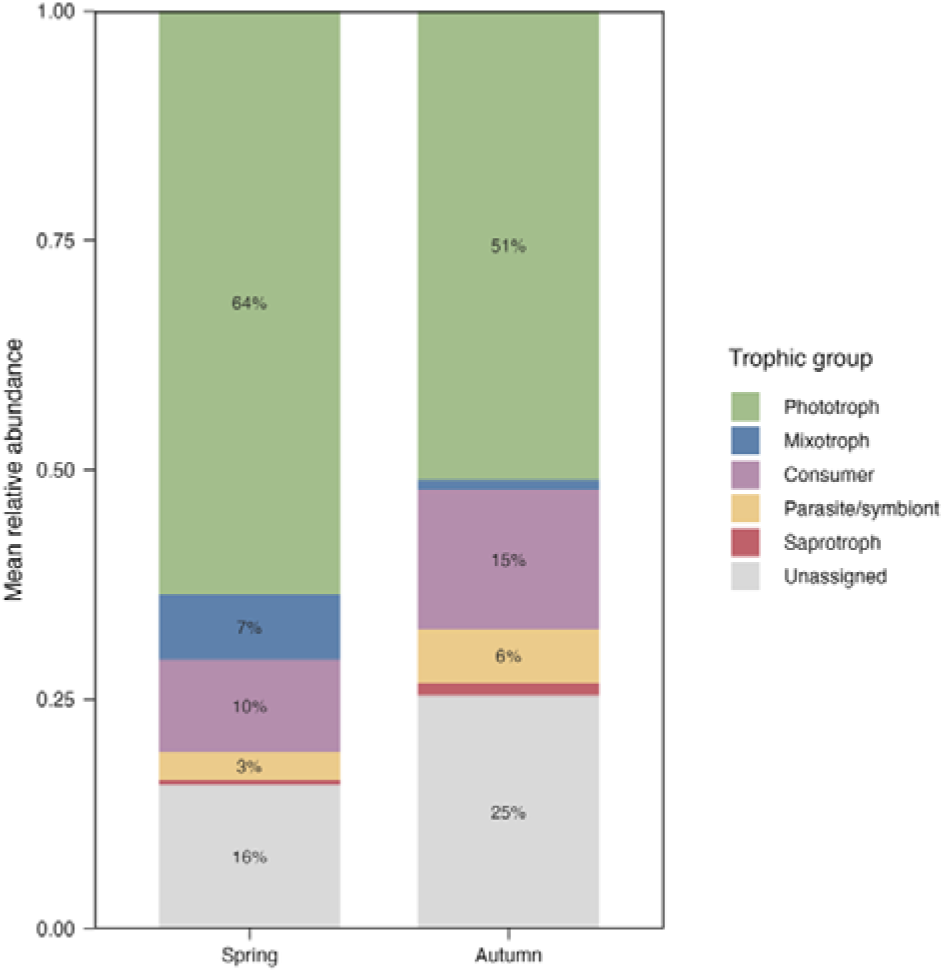
Mean relative abundance of microeukaryotic trophic groups in river biofilms in spring and autumn. Values are shown for groups contributing >3% of reads.

**Figure S2.**
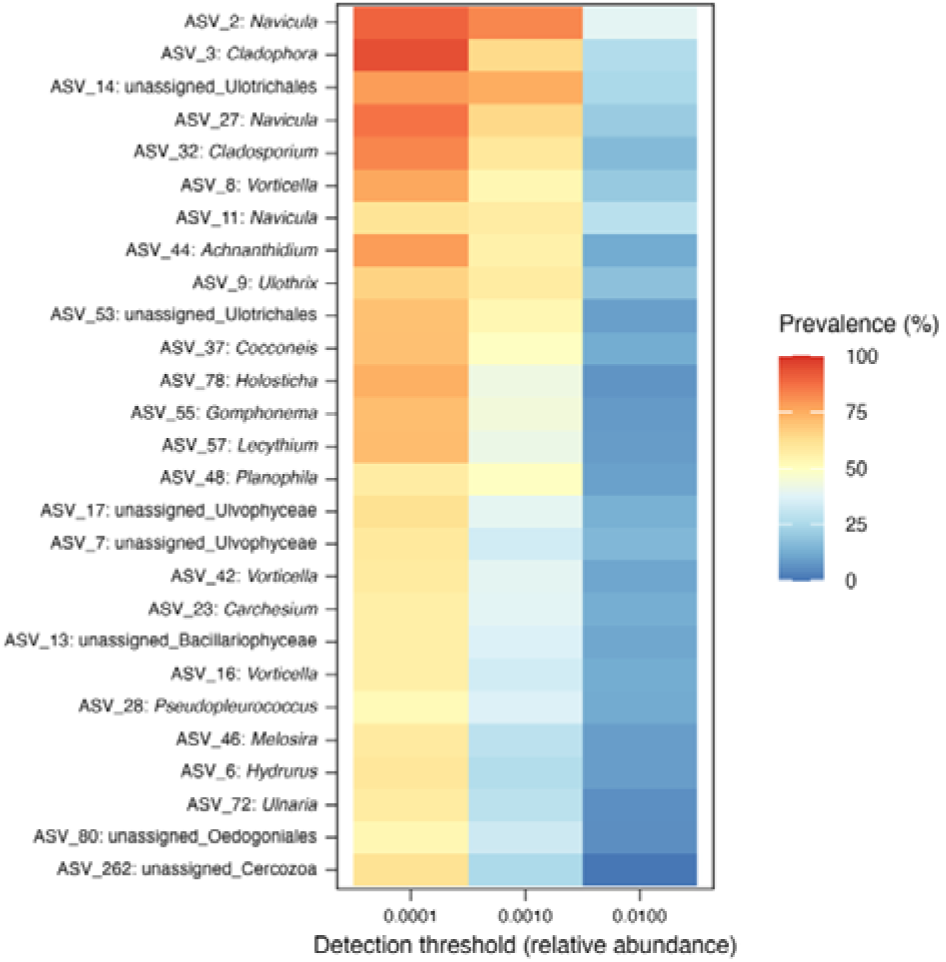
Relative abundance and prevalence of the core community of ASVs, defined by a relative abundance >0.0001 in >50% of samples.

**Figure S3.**
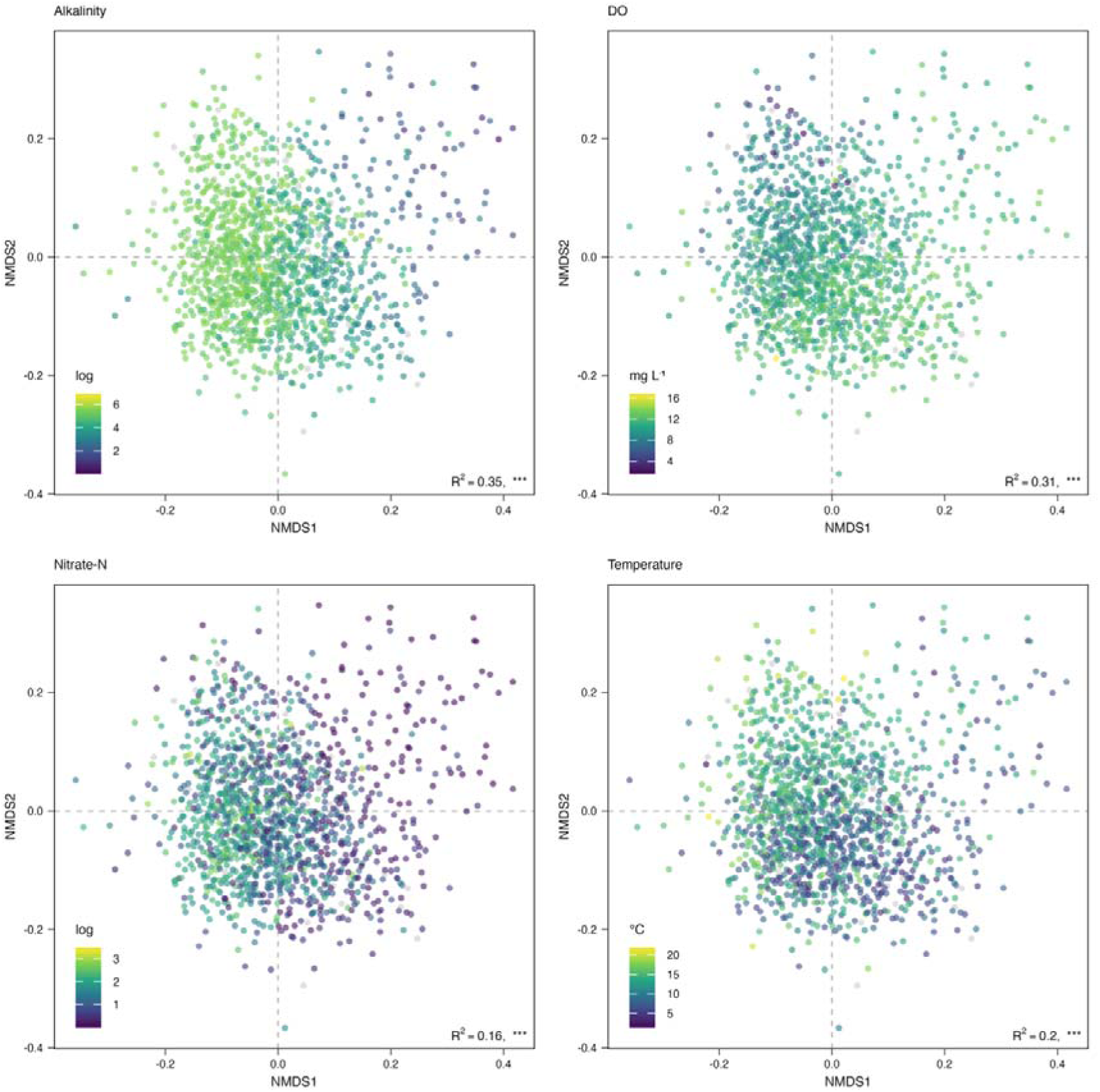
NMDS where samples are coloured by gradients of selected water chemistry variables: (a) alkalinity, (b) DO, (c) nitrate-N, and (d) temperature. NA values are shown in grey. Alkalinity and nitrate-N were visualised on a logarithmic scale. R^2^ and significance levels are shown, p < 0.001 = ***.

**Figure S4.**
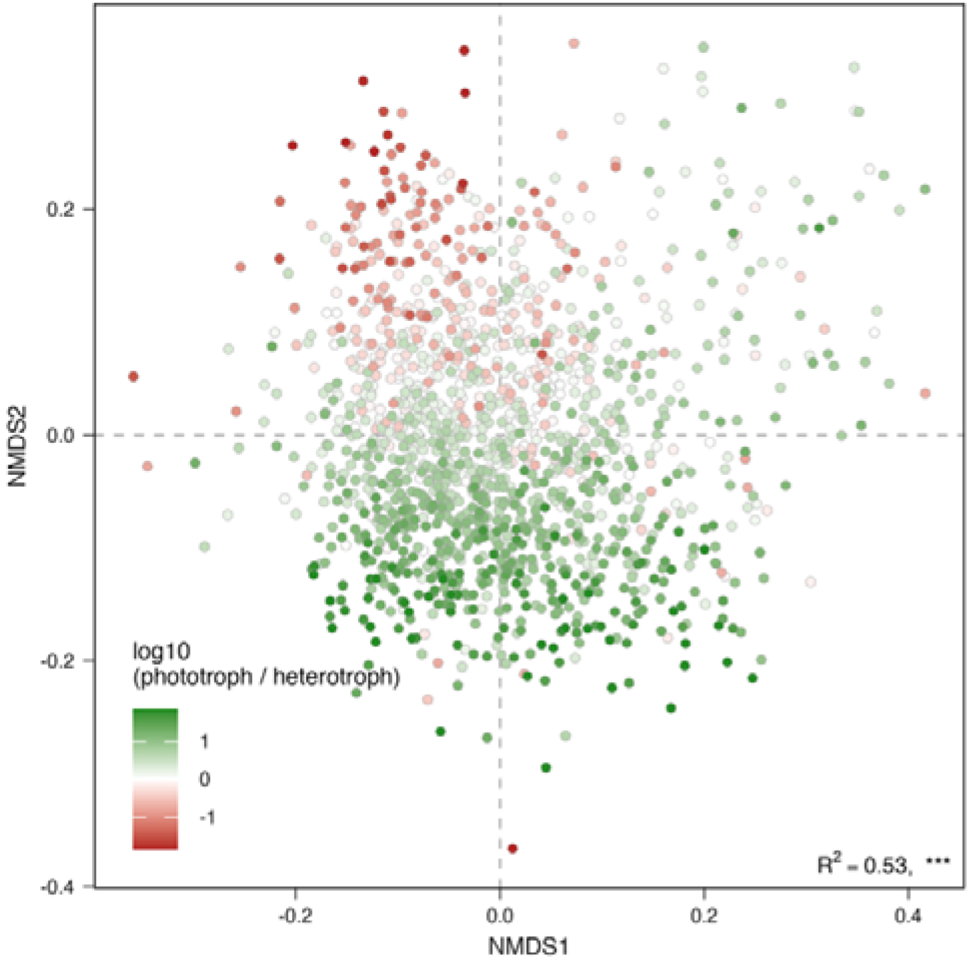
NMDS with points coloured by the log_10_ ratio of phototrophs and heterotrophs, where positive values indicate phototroph-dominated communities, and negative values indicate heterotroph-dominated communities.

**Table S1.**
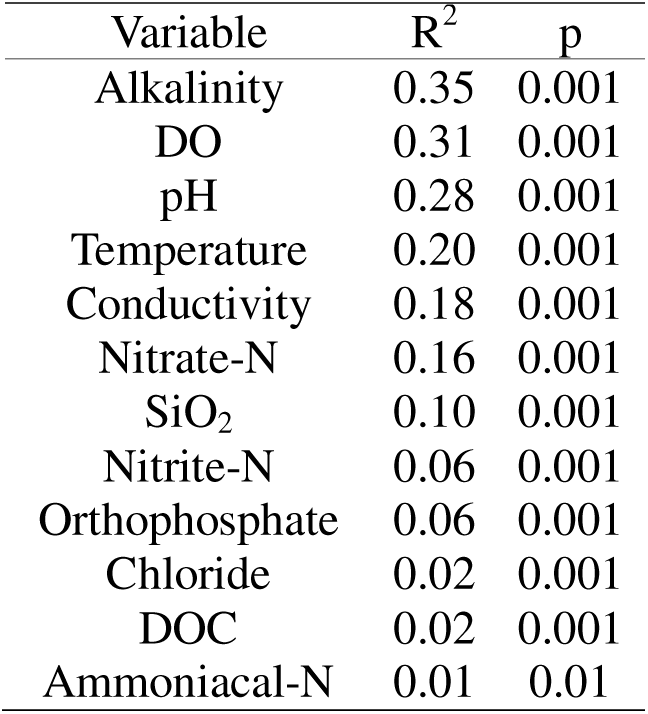
Results of the permutation test between the beta diversity Bray-Curtis dissimilarity matrix and water chemistry variables.

**Table S2.** Mantel and partial Mantel tests of the correlation between community dissimilarity and geographic distance and water chemistry distance, run separately for each survey (season and year). Partial Mantel tests show the correlation with one distance matrix while controlling for the other (e.g., geographic | water chemistry is geographic distance controlling for water chemistry). Values are Mantel r with p values in parentheses, and n denotes the number of samples in each survey.

| Season | Year | n | Mantel |  | Partial Mantel |  |
|---|---|---|---|---|---|---|
|  |  |  | Geographic | Water chemistry | Geographic \| water chemistry | Water chemistry \| geographic |
| Spring | 2021 | 275 | 0.16 (0.001) | 0.40 (0.001) | 0.10 (0.001) | 0.39 (0.001) |
|  | 2022 | 227 | 0.20 (0.001) | 0.48 (0.001) | 0.11 (0.001) | 0.45 (0.001) |
|  | 2023 | 138 | 0.11 (0.001) | 0.45 (0.001) | 0.05 (0.059) | 0.44 (0.001) |
| Autumn | 2021 | 263 | 0.16 (0.001) | 0.38 (0.001) | 0.11 (0.001) | 0.36 (0.001) |
|  | 2022 | 189 | 0.13 (0.001) | 0.38 (0.001) | 0.08 (0.001) | 0.36 (0.001) |
|  | 2023 | 159 | 0.10 (0.002) | 0.44 (0.001) | 0.02 (0.199) | 0.44 (0.001) |

**Figure S6.**
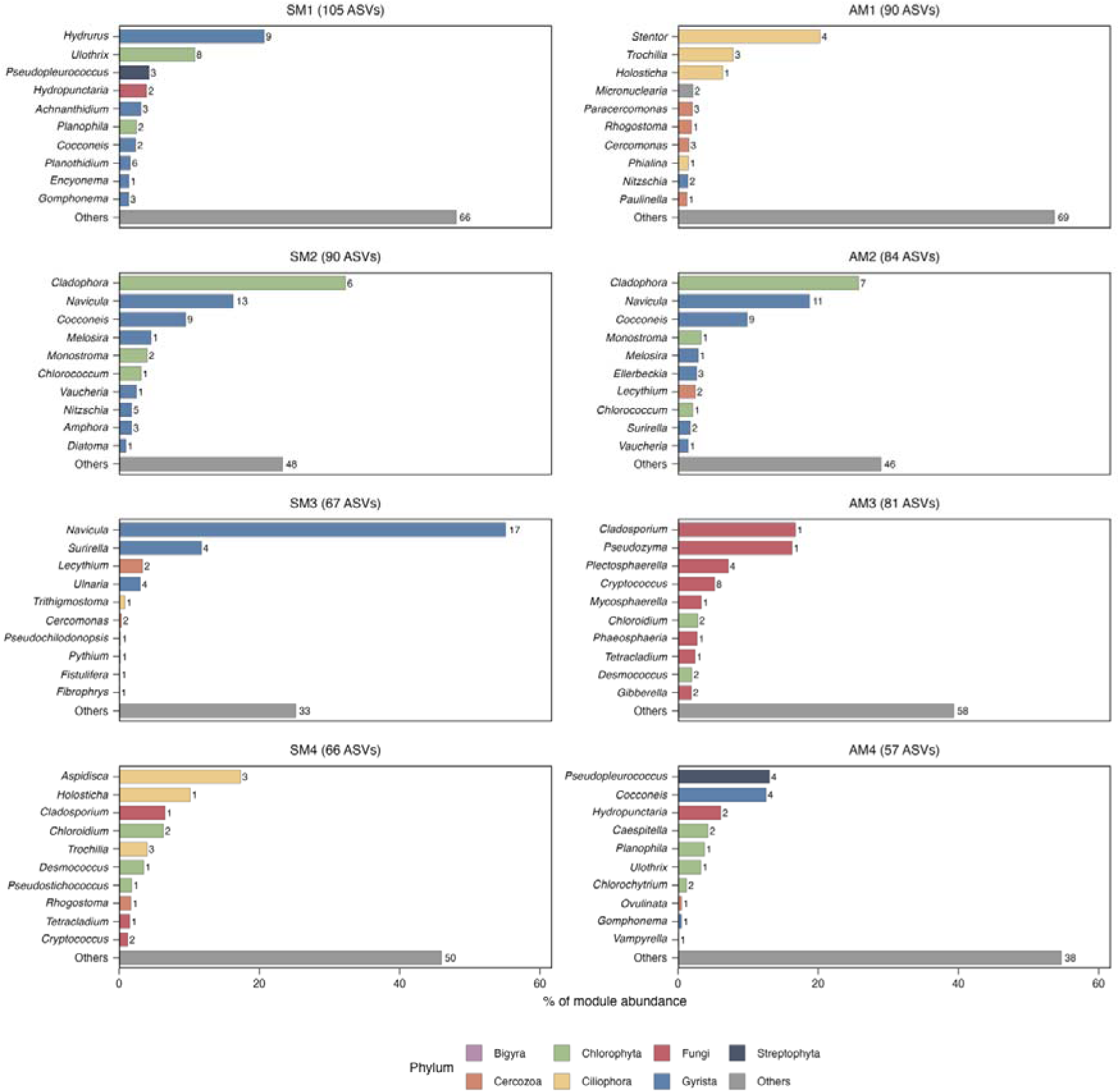
Genus composition of the four largest co-occurrence modules in spring (SM1-4) and autumn (AM1-4). Bars show each genus’s contribution to module abundance measured as the summed relative abundance of its ASVs as a percentage of total module abundance. The ten genera contributing most to each module are shown, and the remaining genera and ASVs unassigned at genus level are grouped as ‘Others’. Bars are coloured by phylum, and numbers give the number of ASVs per genus present in the module.

**Figure S7.**
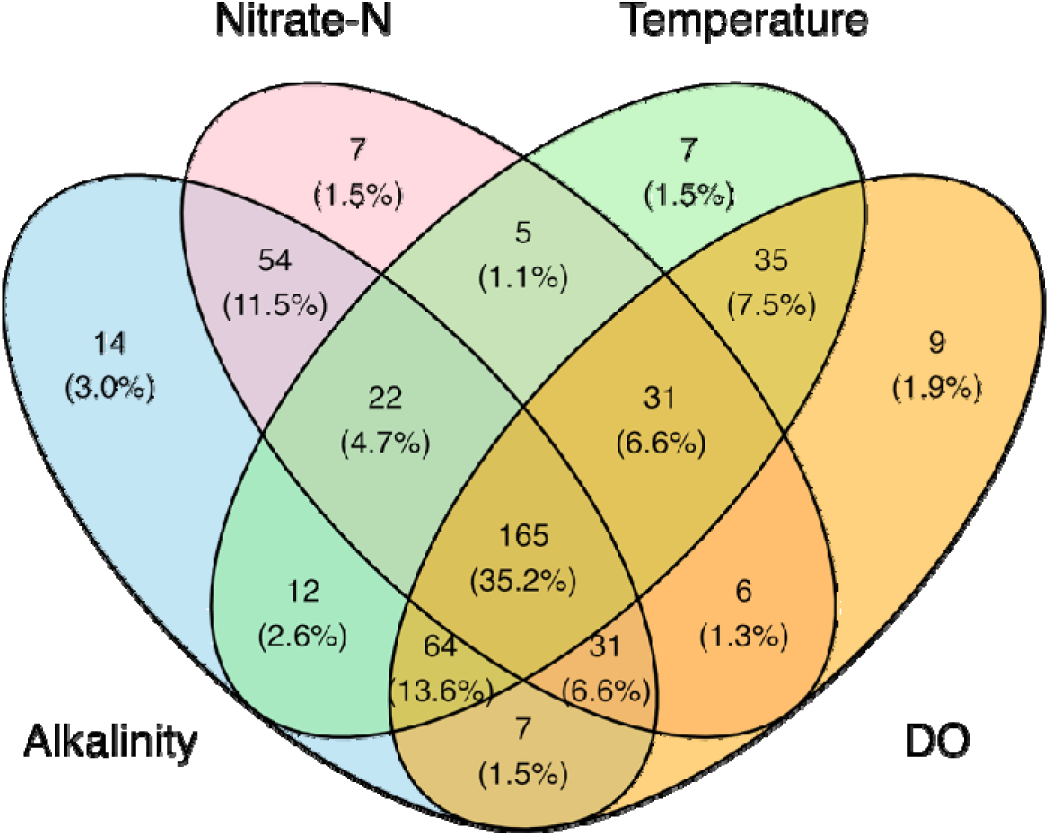
Number and proportion of shared and unique ASVs identified with TITAN as pure and reliable indicators of alkalinity, nitrate-N, temperature, and DO.

